# Independent erosion of conserved transcription factor binding sites points to shared hindlimb, vision, and scrotum loss in different mammals

**DOI:** 10.1101/197756

**Authors:** Mark J. Berger, Aaron M. Wenger, Harendra Guturu, Gill Bejerano

## Abstract

Genetic variation in *cis*-regulatory elements is thought to be a major driving force in morphological and physiological change. However, identifying transcription factor binding events which code for complex traits remains a challenge, motivating novel means of detecting putatively important binding events. Using a curated set of 1,154 high-quality transcription factor motifs, we demonstrate that independently eroded binding sites are enriched for independently lost traits in three distinct pairs of placental mammals. We show that these independently eroded events pinpoint the loss of hindlimbs in dolphin and manatee, degradation of vision in naked mole-rat and star-nosed mole, and the loss of scrotum in white rhinoceros and Weddell seal. Our study exhibits a novel methodology to detect *cis-*regulatory mutations which help explain a portion of the molecular mechanism underlying complex trait formation and loss.

**Author Summary:** Evolution has produced an astounding variety of species with incredibly diverse phenotypes. A central question in evolutionary developmental biology is how (and which) DNA evolves to encode all of these different traits. A prevailing hypothesis is that changes in regulatory DNA, short stretches of DNA which control the expression of protein-coding genes, drive important differences in trait formation between species. The basic building block of regulatory DNA is thought to be transcription factor binding sites, shortl genomic sequences which attract proteins whose central role is to control the rate of transcription. In this study, we asked whether the independent erosion of otherwise highly conserved transcription factor binding sites points to a trait shared between species which have undergone similar adaptations. We show that our method is able to point to the loss of hindlimbs in dolphin and manatee, poor vision in naked mole-rat and star-nosed mole, and loss of scrotum in Weddell seal and white rhinoceros. Overall, our study exhibits a means of detecting evolutionarily important genomic regions which help explain a portion of complex trait loss and retention.

## Introduction

Genetic variation in *cis*-regulatory architecture is thought to be a major force in the morphological and physiological divergence of species(1–3). Although this hypothesis was initially controversial, growing experimental evidence suggests that gene regulation plays a key role in determining phenotypic traits(4,5) such as pelvic reduction in three-spined stickleback(6), loss of wing pigmentation in Drosophila(7), and loss of penile spines in human(8). These observations motivate studying the biological function of transcription factors, their target genes in different cellular contexts, and the regulatory elements to which they bind(9).

While recent studies have examined genome-wide patterns of evolution for transcription factor binding sites(10–12), assigning phenotypic traits to individual sites remains challenging. Despite the conserved sequence-binding preferences of transcription factors, binding sites themselves can erode rapidly over the course of evolution(10,11). On average, individual binding sites are less conserved than the average protein coding gene(13,14) or enhancer(14). Additionally, binding site erosion does not appear to be offset by the creation of new binding events for the same transcription factor. For example, in the case of *CEBPA* and *HNF4A* binding events across five vertebrates, only half of the binding events lost in one lineage appear to be compensated by the creation of another binding event even within 10kb of the original site(11). This rapid turnover of sites, along with the sheer number of binding events and lack of a clear genetic code, make it particularly difficult to ascribe phenotypic function to transcription factor binding events(15).

Despite their rapid turnover, individual binding sites (like individual amino acids) are known to be important building blocks in the formation of complex traits(16), motivating novel methodology of detecting putatively important binding events. Previous work has shown that independent loss of the same complex trait offers a great opportunity to find some of the genomic elements underlying complex trait formation(17,18). Based on those results, we were curious whether independently eroded highly conserved binding sites could also point to an independently lost trait. Conceptually, trait loss in independent clades should lead to <u>neutral</u> drift of all dedicated trait-encoding regions, regardless of the initial inactivating event(18). With sufficient evolutionary time, neutral erosion of these elements becomes detectable between species in which the trait differs. Therefore, in theory, eroded binding sites should congregate in genomic regions with a shared function relevant to an independent trait loss (Figure 1).

**Figure 1.**
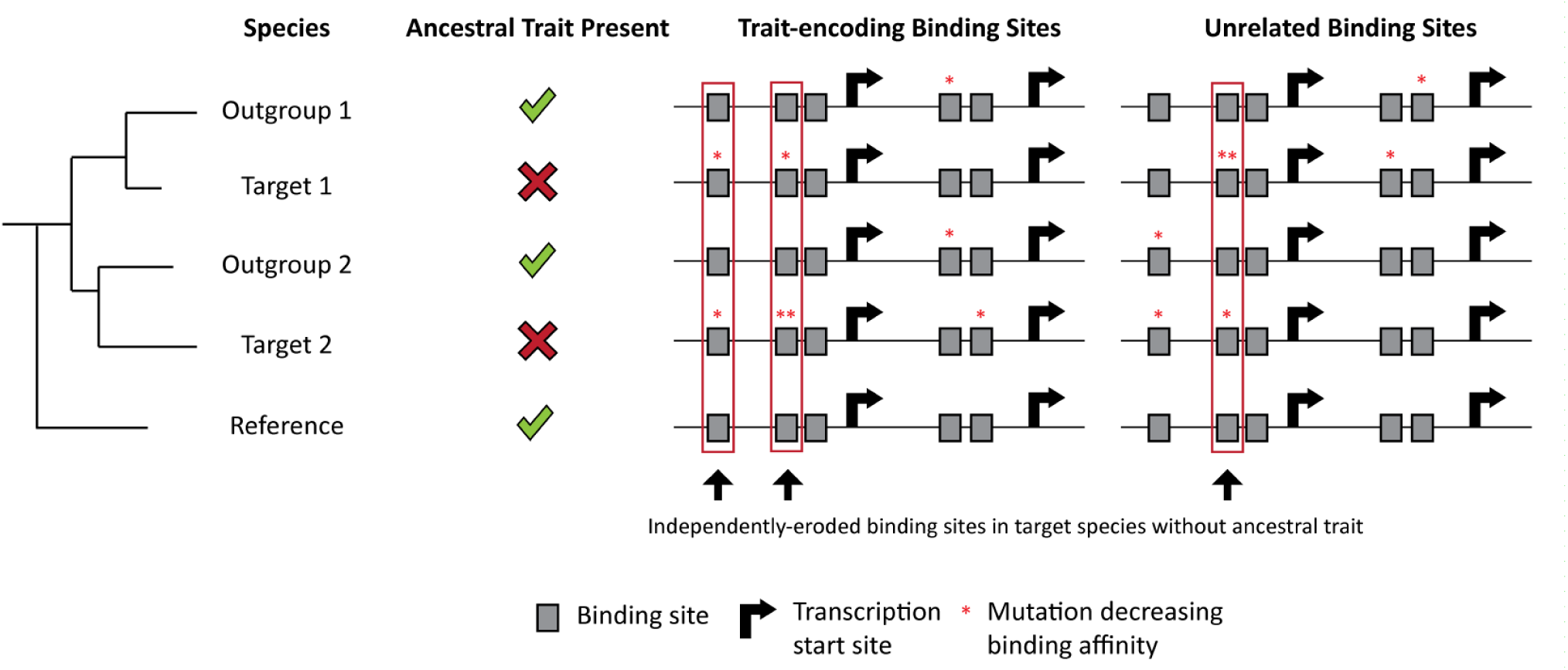
Given enough evolutionary time, independently eroded binding sites should congregate in genomic regions which encode for independently lost traits. As species diverge, a phenotypic trait and the genomic regions required for that trait are inherited from the ancestral species. The trait of interest is necessary to maintain fitness, and therefore important trait-encoding transcription factor binding sites are under negative selection. As target species 1 evolves, a trait-loss event fixates within the species. However, the sister species outgroup 1 still maintains the trait. Since the trait is lost in target species 1, all trait-dedicated information now switches to neutral selection in the species. This leads to neutral erosion of trait-encoding transcription factor binding sites. Similarly, in target species 2, a trait-loss event (but not necessarily the same event as in target species 1) for the same trait fixates in the population.Here too, sister species outgroup 2 still maintains the trait. Now all trait-encoding information in target species 2 switch to neutral selection, and therefore the trait-encoding binding sites begin to neutrally erode. Using the sister species as outgroups, we can identify all transcription factor binding sites which have eroded in our target species but have been maintained in our outgroup species and many other references species. We refer to these sites as independently eroded binding sites. This very unusual evolutionary signature is shown in Figure 2 and Table 1 to be strongest next to key genes for the development of an important independently lost complex trait.

Using a large set of transcription factor binding site motifs, we set out to test this model by identifying binding sites which are eroded in two independent clades, but are otherwise highly conserved across the phylogeny of placental mammals. For each pair of species, we performed a statistical test with 3,538 ontology terms from the Mouse Genome Informatics (MGI) Gene Expression Database(19) to determine the most significant shared biological function of the independently eroded transcription factor binding sites. Using our novel test, we recapitulate three diverse loss-of-function phenotypes over distinct clades, suggesting the generality of the approach.

## Results

### Loss of hindlimbs in aquatic mammals

Cetacean and Sirenian lineages have independently adapted to an aquatic habitat from their terrestrial ancestral species(20) (Figure 2A). Using the dolphin and manatee genomes as representative of each respective clade, we asked whether independently eroded binding sites in dolphin and manatee are associated with any morphological divergence of aquatic mammals.

**Figure 2.**
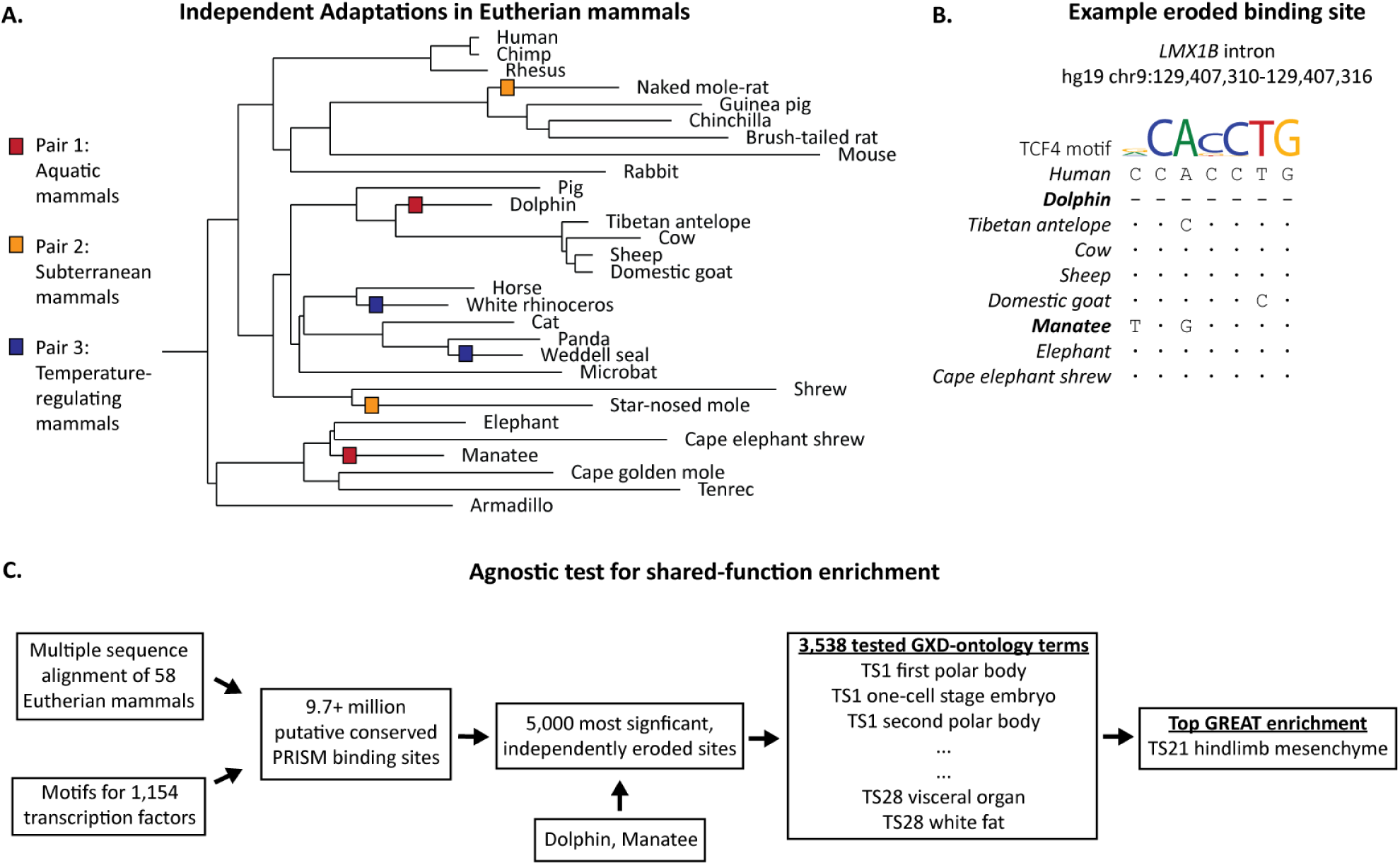
Independently eroded binding sites are enriched next to key genes for independent complex trait losses. (A) Throughout the evolution of placental mammals, species have independently adapted to a variety of environments. In the three pairs of species shown here, we test whether independently eroded binding sites are statistically enriched for functions associated with these adaptations (see Supplementary Figure 1 for the phylogeny of all mammals used in this study, and Figure 1 for the rationale). (B) Independently eroded binding sites can either be the result of a deletion (i.e. dolphin in this example) or mutations which decrease binding affinity (i.e. manatee). Bases identical to human are represented as dots and single dashes represent deleted bases. (C) Using a library of 1,154 transcription factor motifs, we identify approximately 9.7 million putative mammalian conserved binding sites. We consider a conserved site to be eroded in the target species if a motif match is absent in the target species, but present in any of the outgroup species. Binding sites are considered independently eroded if they are eroded along two distinct clades of species. For a given pair of species, the 5,000 most significant independently eroded sites are agnostically tested against 3,538 ontology terms from the MGI Gene Expression Database (GXD) to identify a most significant shared function (see Table 1).

To perform binding site prediction, we obtained an hg19-anchored MULTIZ alignment of 58 Eutherian mammals from the UCSC Genome Browser(21) (Supplementary Table 1). We also curated a set of 1,154 high-quality transcription factor motifs from UniProbe(22), JASPAR(23), and TransFac(24). Both monomeric and dimeric motifs were included because previous work has demonstrated that complexes have modified binding affinities(25,26). With these resources, we used PRISM(27) to identify binding sites which are much more conserved than their surrounding sequence, suggesting evolutionary constraint and functional importance of the sites (see Methods). This method has been shown to substantially outperform conservation free position weight matrix prediction, as well as generate transcription factor binding profiles which are more similar to ChIP-seq data(27). In total, we identified 9,729,644 putative conserved transcription factor binding sites covering 0.848% of the reference genome (Figure 2C).

Eroded transcription factor binding sites were defined as binding sites predicted in at least one outgroup species, but not in the target species of interest. For dolphin, the following species are present in the multiple alignment as the outgroup: Tibetan antelope, cow, sheep, and domestic goat. For manatee, the multiple alignment offers elephant and cape elephant shrew as the outgroup species (Figure 2A-B). We discarded any eroded binding site prediction that overlapped an assembly gap in the target species(18). We identified 826,796 and 703,642 putative eroded binding sites in dolphin and manatee, respectively, with the intersection containing 80,798 independently eroded binding sites shared by both target species.

To assess the shared function of the most surprising erosion events, we selected the top 5,000 most highly conserved sites (plus 11 ties at the cutoff point), according to PRISM (Supplementary Table 2). These binding sites were supplied to GREAT (Genomic Regions Enrichment of Annotations Tool)(28) to agnostically determine the most significant shared function among the sites. Using GREAT’s original optimized settings, our analysis was conducted over 3,538 ontology terms from the MGI Gene Expression Database(19), a compendium of mouse developmental time points (Theiler stage, or TS1-28) with their corresponding validated expressed genes. The significance thresholds optimized in the original GREAT paper were used to define statistical significance: a region-based fold enrichment of at least 2 and a false-discovery rate (FDR) of less than or equal to 0.05 for both the region-based and gene-based tests.

Strikingly, using these criteria, the most enriched term for the set was “TS21 hindlimb mesenchyme” with a region-based FDR of 1.3 x 10^-4^ and a region-based fold enrichment of 2.85 (Table 1). Indeed, both clades have experienced a dramatic reduction or loss of hindlimbs(20,29). It is thought that the loss of hindlimbs contributes to streamlining the body, reducing drag and therefore drastically reducing the amount of energy necessary for swimming(30). Our results identify a portion of the molecular signature underpinning the shared feature and its subsequent loss.

**Table 1.**
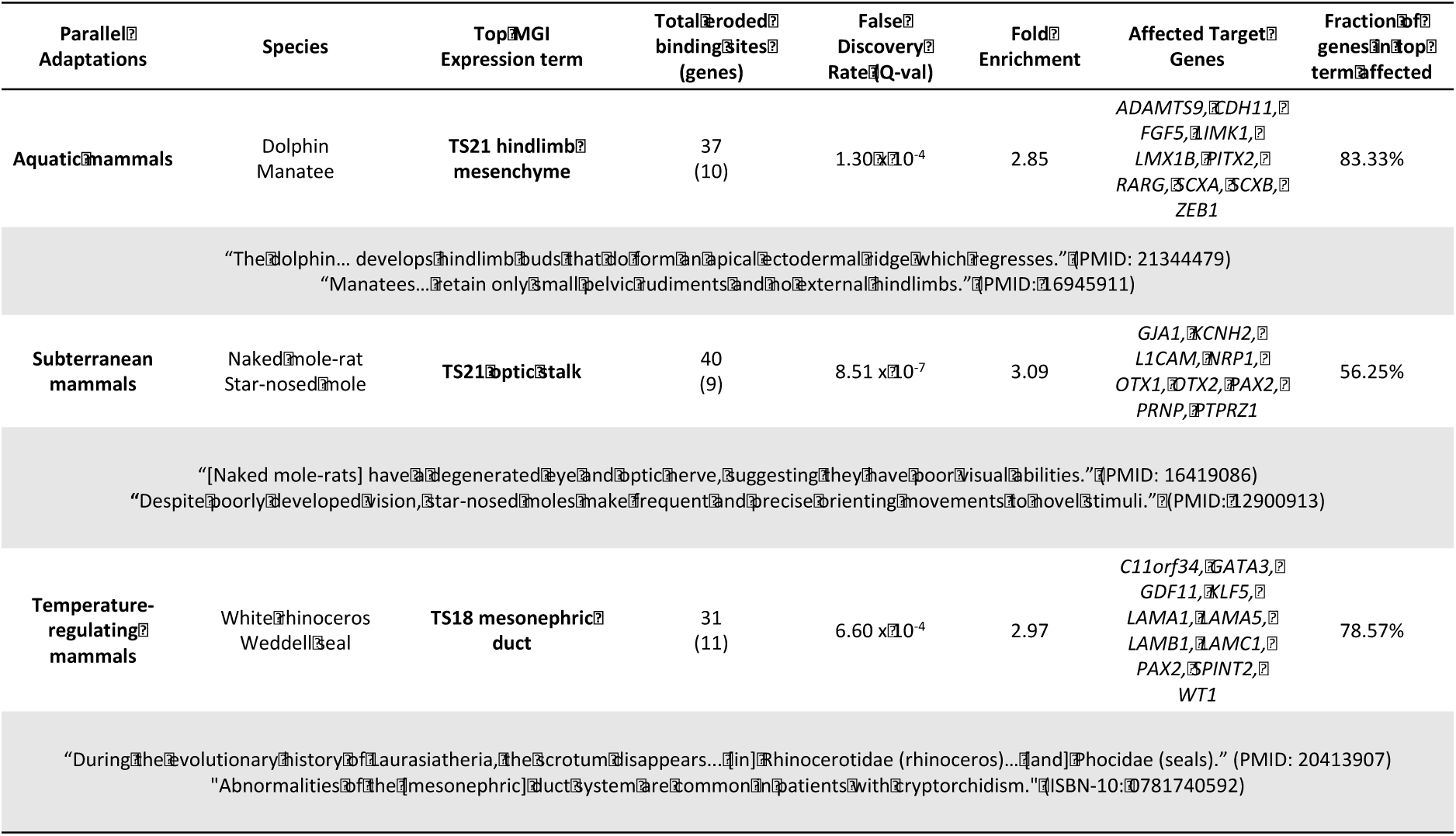
Independently eroded binding sites congregate in the regulatory domain of important trait relevant genes. See Figures 1-2 for the test we perform. TS – Theiler Stage.

### Degradation of vision in subterranean mammals

Both naked mole-rat and star-nosed mole have independently adapted to a subterranean habitat (Figure 2A). Using naked mole-rat and star-nosed mole as our target species, we asked whether independently eroded binding sites are associated with any morphological divergence of subterranean mammals.

For naked mole-rat, the following species were available as an outgroup: Guinea pig, chinchilla, and brush-tailed rat. For the star-nosed mole, shrew served as the outgroup species (Figure 2A). Using the exact same methods from the aquatic mammals’ analysis, we identified 969,015 and 797,304 putative eroded binding sites in naked mole-rat and star-nosed mole, respectively, with the intersection resulting in 105,416 independently eroded binding sites. Again, we selected the top 5,000 most significant sites plus 121 ties (Supplementary Table 3). Using GREAT, the most enriched term for this set was “TS21 optic stalk”, a precursor structure to the optic nerve(31), with a region-based FDR of 8.51 x 10^-7^ and a region-based fold enrichment of 3.09 (Table 1). Indeed, adaptation to a subterranean environment has led to regressive evolution of the visual system, with key vision-related genes eroded in these species(32,33).

### Loss of scrotum in mammals with temperature-regulating adaptations

Both phocid seals and rhinoceroses (Figure 2A) experience a wide range of external temperatures in their natural habitat. Both of these clades are able to maintain a relatively low body temperature compared to other Artiodactyla species(34), piquing interest in how both clades regulate their core body temperature(35,36). Using white rhinoceros and Weddell seal as the target species, we asked whether independently eroded binding sites are associated with any morphological divergence common to these temperature-regulating species.

For white rhinoceros, horse was available as the outgroup species. For Weddell seal, panda served as the outgroup species (Figure 2A). Using the previously described methods, we identified 447,297 and 425,706 putative eroded binding sites in white rhinoceros and Weddell seal, respectively, with the intersection resulting in 31,097 independently eroded binding sites. We selected the top 5,000 most significant sites plus 102 ties (Supplementary Table 4). Using GREAT, the most enriched term for this set was “TS18 mesonephric duct” with a region-based FDR of 6.6 x 10^-4^ and a region-based fold enrichment of 2.97 (Table 1). The mesonephric duct is an embryonic tissue which forms the epididymis and vas deferens, both of which are commonly malformed in patients with cryptorchidism, the absence of testes from the scrotum(37). Indeed, both white rhinoceros and Weddell seal do not form a scrotum(38). Previous work speculates that the loss of the scrotum is in accordance with the development of an intra-abdominal gonadal cooling system in order to optimize sperm production(38).

## Discussion

The interpretation of *cis*-regulatory elements is a difficult task. Our inability to properly identify regulatory elements of interest inhibits the understanding of gene regulation in both evolution and disease. Here, we present a method to computationally identify highly conserved putative binding sites which are independently eroded in two clades of Eutherian mammals, and by extension are dedicated to trait encoding in the species that preserve both trait and sites. We show that for aquatic, subterranean, and temperature-regulating mammals, the erosion of the highly conserved transcription factor binding sites respectively point to the following independent trait losses: loss of hindlimbs, degradation of vision, and loss of scrotum. All tests are computed across 3,538 ontology terms, demonstrating the statistical strength of our most-enriched terms.

Our previous work linked independent erosion of protein-coding genes(18) and highly conserved enhancers(17) to independently eroded phenotypes. While these methods successfully identify genomic regions of large effects, current work suggests that complex traits are shaped by the accumulation of genomic events with small effect sizes(3,39). Therefore, studying the adaption of *cis*-regulatory regions at the resolution of individual transcription factor binding events is necessary to further our understanding of complex trait formation(40). While many binding events are rapidly gained and lost throughout the course of evolution(10,11), our test is able to successfully point to three diverse morphological changes, suggesting that analyses at the resolution of key individual binding sites are surprisingly tractable, likely not only across mammals but across many different phyla.

Furthermore, our test recapitulates genes which are experimentally validated to be involved in the trait losses of interest. For aquatic mammals, the most enriched gene set includes *PITX2*, a homeobox transcription factor required for the formation of hindlimbs(41), and *LMX1B* (Figure 3A). *LMX1B* is a LIM homeodomain transcription factor responsible for dorsal cell fate in the developing limb(42,43). For subterranean mammals, the most enriched ontology term includes *PAX2*, a transcription factor necessary for the development of the optic stalk(44). Rare germline mutations in *PAX2* are known to cause renal colomba syndrome, an autosomal-dominant disease characterized by optic nerve dysplasia(45). The most enriched term also includes *NRP1*, a transmembrane receptor necessary for proper angiogenesis and arteriogenesis of the retina (46) (Figure 3B). Finally, for mammals with advanced temperature regulating capacity, the most enriched ontology term includes *WT1*, a transcription factor involved in both renal and gonadal development (Figure 3C). Patients with rare germline mutations in *WT1* have presented with cryptorchidism, the absence of testes from the scrotum(47). Previous work has also verified the association between *Wt1* and cryptorchidism using conditional gene inactivation mice(47).

**Figure 3.**
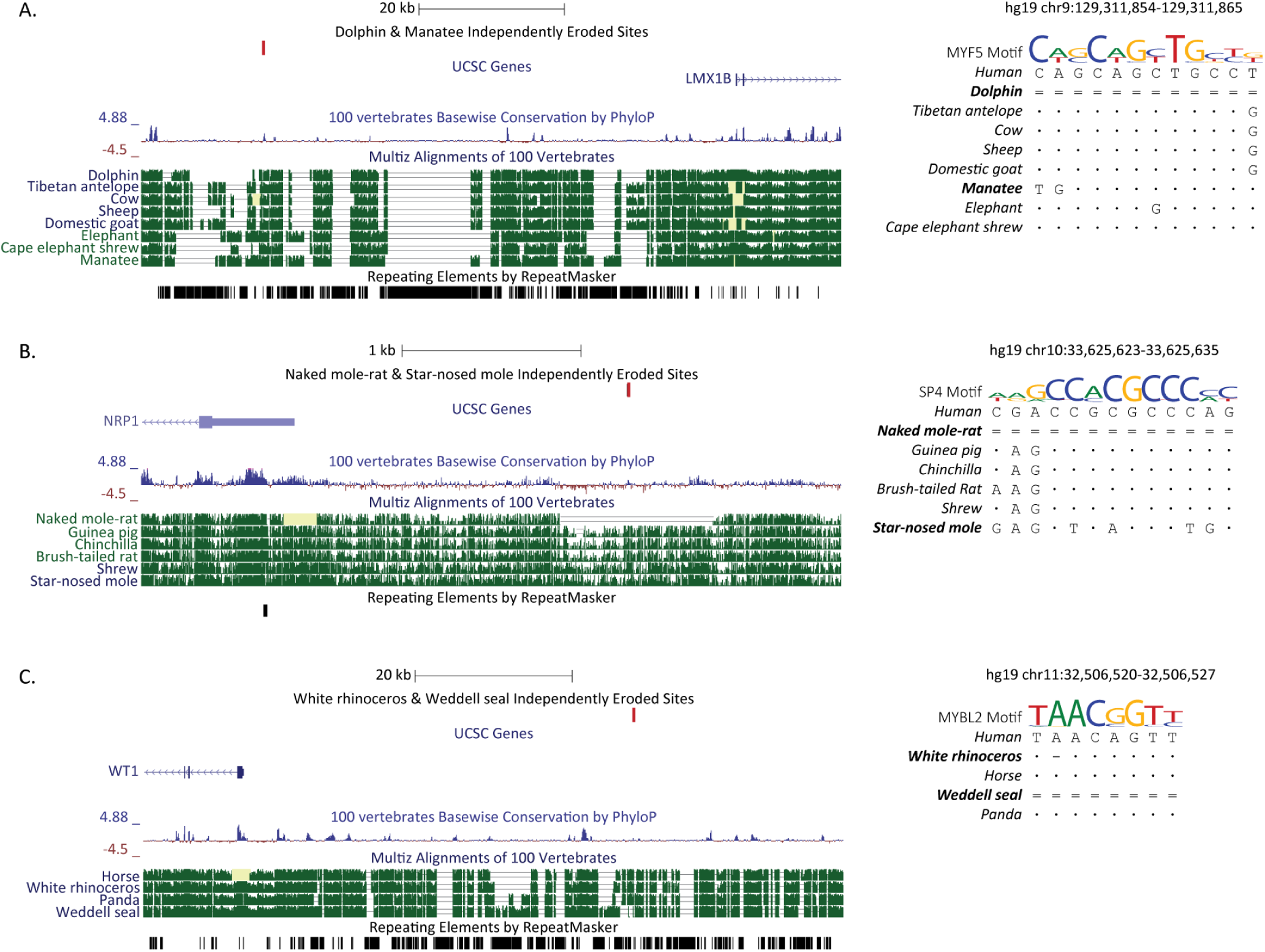
Examples of independently eroded binding sites next to important genes for independent complex trait loss. Bases identical to human are represented as dots, single dashes represent deletions, and double dashes represent non-aligning bases at the query species. (A) In both dolphin and manatee we find an independently eroded *MYF5* binding motif upstream of *LMX1B*. *MYF5* is a myogenic transcription factor, the first myogenic factor to become active in the developing limb bud(51). *LMX1B* is a LIM homeodomain transcription factor responsible for dorsal cell fate in the developing limb(42,43). (B) In both naked mole-rat and star-nosed mole, we find an independently eroded *SP4* binding motif upstream of *NRP1*. *SP4* is a transcription factor which controls transcription of photoreceptor-specific genes in conjunction with *CRX*(52). *NRP1* is a transmembrane receptor necessary for proper angiogenesis and arteriogenesis of the retina (46). (C) In both white rhinoceros and Weddell seal we find an independently eroded *MYBL2* binding motif upstream of *WT1*. *WT1* is a transcription factor involved in both renal and gonadal development. Conditional inactivation of *Wt1* in mice causes left-sided cryptorchidism with 40% penetrance(47).

ChIP-seq, along with efforts such as the ENCODE Project(48), have transformed our ability to observe transcription factor binding. Despite these extensive resources, determining the phenotypic contribution of binding events is difficult. By identifying independently eroded binding sites and demonstrating that these sites point to independently lost morphological traits, we identify a small set of putative binding sites primed for experimental follow-up. As the research community sequences more genomes, characterizes the binding motifs of additional transcriptions factors, and improves methods to model transcription factor binding, the ability to identify interesting binding events with our method will continue to improve. Coupled with the accuracy of CRIPSR/Cas genome-editing systems, our method enables the dissection of phenotypically relevant *cis*-regulatory regions to further biological understanding of complex-trait formation.

## Materials and Methods

### Multiple genome alignment and phylogenic tree

A hg19-anchored MULTIZ alignment of 100 species, along with the corresponding phylogenetic tree and branch lengths, was obtained from the UCSC Genome Browser(21) and subset to the provided 58 placental mammals (Supplementary Table 1, Supplementary Figure 1). All analyses were performed on the MULTIZ alignment of 58 mammals. Exon and repeat annotations were obtained from the UCSC Genes and RepeatMasker tracks, respectively.

### Transcription factor binding motif library

The set of transcription factor binding motifs consists of 1,154 unique high-quality monomer and dimer motifs for 753 transcription factors. Motifs were curated from existing databases and primarily literature as previously described(40). Both monomers and dimer motifs were included because previous studies have demonstrated that complexes have modified binding affinities(25,26).

### Identifying Eroded Transcription Factor Binding Sites

Binding site prediction was performed on the multiple alignment of 58 Eutherian mammals using PRISM (27). PRISM performs binding site prediction by searching for motif-matching sequences which are both conserved across multiple species and more conserved than their surrounding sequence, suggesting evolutionary constraint on the binding site. To identify these constrained sites, we first partitioned the reference genome into windows of conservation. Each base pair was assigned a weighted percent identity score by calculating the total branch length over which the base pair is conserved normalized by the total branch length of the phylogeny. The percent identity score for each base was then smoothed by averaging the percent identity scores of a 100bp window centered upon the base pair of interest. Conservation values over the entire multiple alignment were then grouped into 1% bins.

Next, for each binding motif, a set of null model motifs was generated by shuffling the columns of the position weight matrix. Adjacent CpG columns were conserved during shuffling. The shuffled motifs were then used to generate a set of predictions across the reference genome. Sequences with a MATCH(49) score of at least 0.8 were considered matches to the binding affinity matrix. For each 1% bin of conservation defined above, we recorded the distribution of Bayesian branch length (BBL) scores(50) for all sequences which match a shuffled motif. Finally, the true motif was used to generate a set of predictions along the reference genome. For each prediction, we derived an empirical p-value based on the distribution of BBL scores for the respective conservation bin. Formally, given the distribution of scores for the respective conservation bin, the empirical p-value is the probability that a shuffled motif score is greater than or equal to the score of the real motif. A prediction was only retained if it had an uncorrected conservation p-value less than or equal to 10^-3^. Additionally, a prediction was only retained if the binding site is preserved in at least five species with a total phylogenetic branch length of 3.0 substitutions per site or more. Predictions were generated for the 58 placental mammals over all regions which did not contain an exon or a repeat element in the reference genome. See the PRISM paper for a more detailed explanation(27).

For each species of interest, one or more outgroup species were dictated by the UCSC alignment as a reference to define a set of lineage-specific eroded sites. We selected the lowest common ancestor which was thought to not exhibit the trait based on observations in extant species. All species derived from the least common ancestor which do not exhibit the trait of interest were used as the outgroup species. We defined the eroded binding sites of a species to be the binding site predictions which are present in any of the respective outgroup species (and therefore likely also in the ancestor), but not in the species of interest. Eroded site predictions overlapping assembly gaps were discarded(18).

### Inferring statistically significant accumulation of eroded binding sites near genes that share a common function

For a pair of target species, the eroded sites in each species were intersected to produce a set of independently eroded sites. These sites were ranked by their excess conservation score, and the top 5,000 sites were used to form a high-confidence set of independently eroded sites. In order to prevent arbitrary tie-breaking, additional predictions were included if their excess conservation p-value was equivalent to the significance of the last prediction in the ranked list. Since paralogous transcription factors often have similar binding motifs, we required all entries in our set of most significant sites to not overlap one another. If multiple predictions did overlap, only the prediction with the highest excess conservation score was retained.

Each set of high-confidence independently eroded sites was submitted to GREAT (Genomic Regions Enrichment of Annotations Tool) v3.0.0 (28). Binding sites were associated with target genes using the default “Basal plus extension” association rule with the default distances of 5kb upstream and 1kb downstream from the canonical transcription start site, with an extension of up to 1Mb or the next gene. The optimized thresholds from the original GREAT paper were used to define statistical significance: a region-based fold enrichment of at least 2, and a false-discovery rate of less than or equal to 0.05 for both the region-based and gene-based tests. Analyses were conducted over the MGI Gene Expression Database(19) for all terms *a priori* annotated with at least 10 genes and at most 500 genes, resulting in 3,538 tests.

## Acknowledgements

We thank J. Birgmeier, H. I. Chen, W. Heavner, K. Jagadeesh, A. Marcovitz, and members of the Bejerano Laboratory for valuable discussions and project feedback. We also thank the UCSC Browser Team for providing the MULTIZ alignment. This work was funded by a Stanford Engineering Excellence and Diversity Fellowship and a National Science Foundation Graduate Research Fellowship DGE-1656518 (MJB), as well as NIH grants U01MH105949 and R01HG008742, a Packard Foundation Fellowship and a Microsoft Faculty Fellowship (GB).

## Author Contributions

Conceived and designed the study: MJB, GB. Performed the experiments: MJB. Analyzed the data: MJB, GB. Contributed materials/analysis tools: AMW, HG. Wrote the paper: MJB, GB. All authors reviewed the manuscript.

## Supplementary Information Legends

**Supplementary Table 1**

**Species List.** The list of 58 placental mammals and their respective genome assemblies used in this study.

**Supplementary Table 2-4**

**Most significant independently eroded transcription factor binding sites.** A full list of the most significant independently eroded binding sites for each pair of species. Genomic coordinates are given relative to the reference genome (human, hg19). Coordinates highlighted in green are binding sites associated with the most enriched term (Table 1). SI Table 2) dolphin and manatee; SI Table 3) naked mole-rat and star nosed-mole; SI Table 4) Weddell seal and white rhinoceros.

**Supplementary Figure 1.**
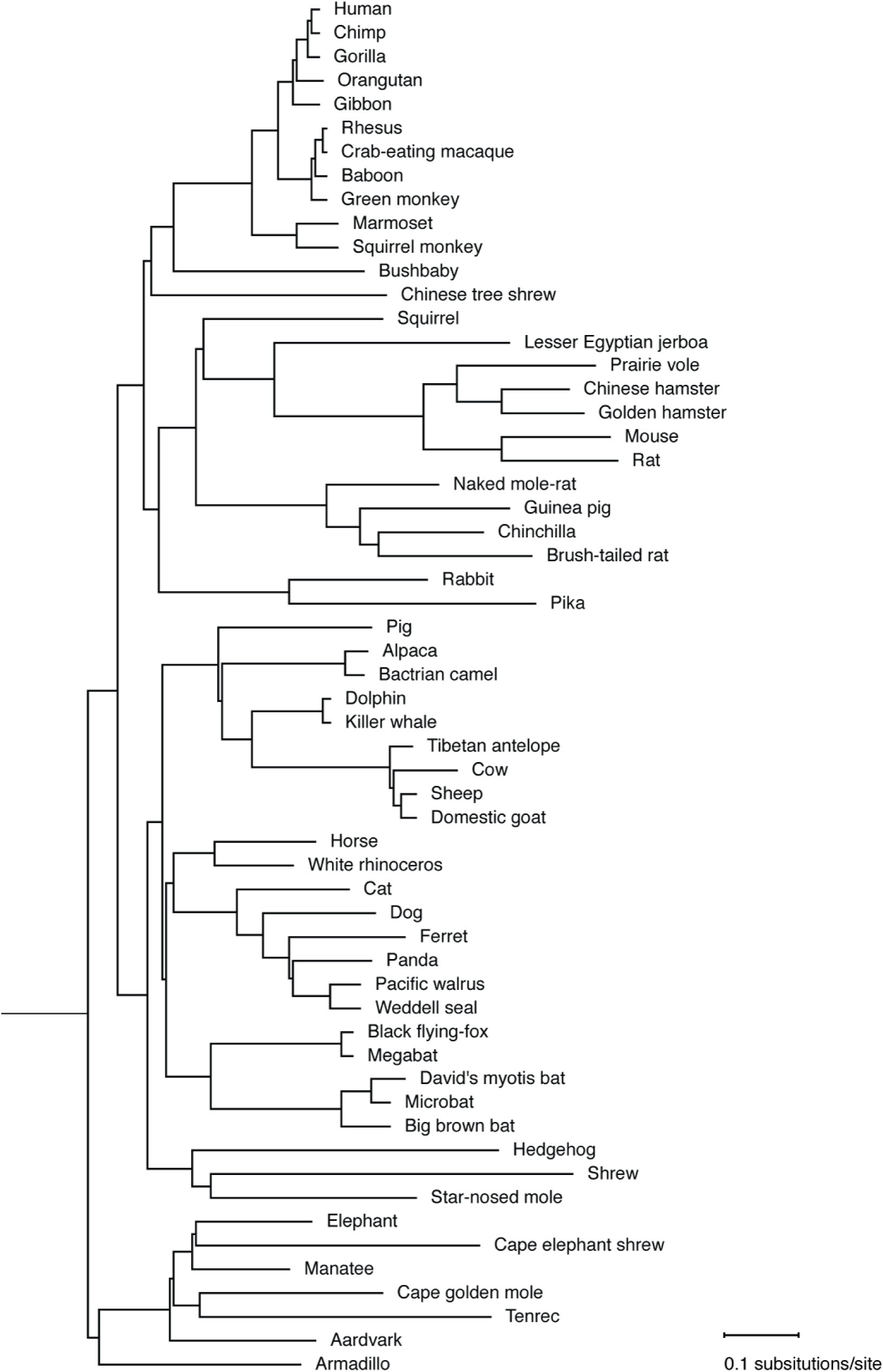
Weighted phylogenetic tree. A UCSC derived substitution per site weighted tree of the 58 species used in this study.

